# Western-style diet impedes colonization and clearance of gut bacterial pathogens

**DOI:** 10.1101/2020.10.06.329144

**Authors:** Jun Zou, Xu Zhao, Juan Noriega, Andrew T. Gewirtz

## Abstract

**Background:** Western-style diet (WSD), which is high in fat and low in fiber, lacks nutrients to support gut microbiota. Consequently, WSD promotes microbiota encroachment and reduces microbiota density, potentially influencing colonization resistance, immune system readiness, and, consequently host defense against pathogenic bacteria. Additionally, the low-nutrient colonic environment resulting from WSD might impact bacterial pathogens.

**Aims:** Examine impact of WSD on infection and colitis in response to gut bacterial pathogens.

**Methods:** Mice fed grain-based chow (GBC), WSD, or various versions thereof, were orally infected with *Citrobacter rodentium* or *C. difficile*. Colonization and its consequences, including inflammation and death were monitored.

**Results:** WSD delayed *Citrobacter* growth, reduced its virulence gene expression and ameliorated inflammation. However, while GBC-fed mice uniformly cleared *Citrobacter* and were impervious to subsequent *Citrobacter* challenges, most WSD-fed mice remained chronically infected with *Citrobacter* while those that cleared it were highly prone to re-infection. Such persistent proneness to *Citrobacter* infection did not reflect reduced immune responsiveness but rather reflected reduced colonization resistance likely from the low microbiota density resulting from WSD feeding. While persistent *Citrobacter* infection did not cause overt inflammation, it was associated with low-grade inflammation in colon and adipose tissue that was associated with insulin resistance. An analogous pattern was seen in response to *C. difficile* with WSD resulting in delayed colonization and mortality while enriching WSD with fiber hastened colonization but afforded clearance and survival.

Altering microbiota via diet can profoundly impact the course and consequence of infection following exposure to gut bacterial pathogens.

**Synopsis:** Dietary fiber results in a nutrient rich colon that is exploited by gut pathogens including *C. rodentium* and *C. difficile*, that rapidly colonize the colon. Hight-fat, low-fiber western-style diet results in a low-density gut microbiota unable to mediate clearance of pathogens from the colon. Inability to clear pathogens from the gut lumen promotes chronic low-grade inflammation and insulin resistance.

**Graphic summary:** The low-fiber content of WSD impedes initial colonization of pathogens. Immune system functions appropriately irrespective of diet to clear pathogens from the mucosa. Commensal microbiota of mice fed GBC but not WSD is sufficient to eliminate pathogen from the lumen of the intestine. Underlying mechanisms discussed in text.

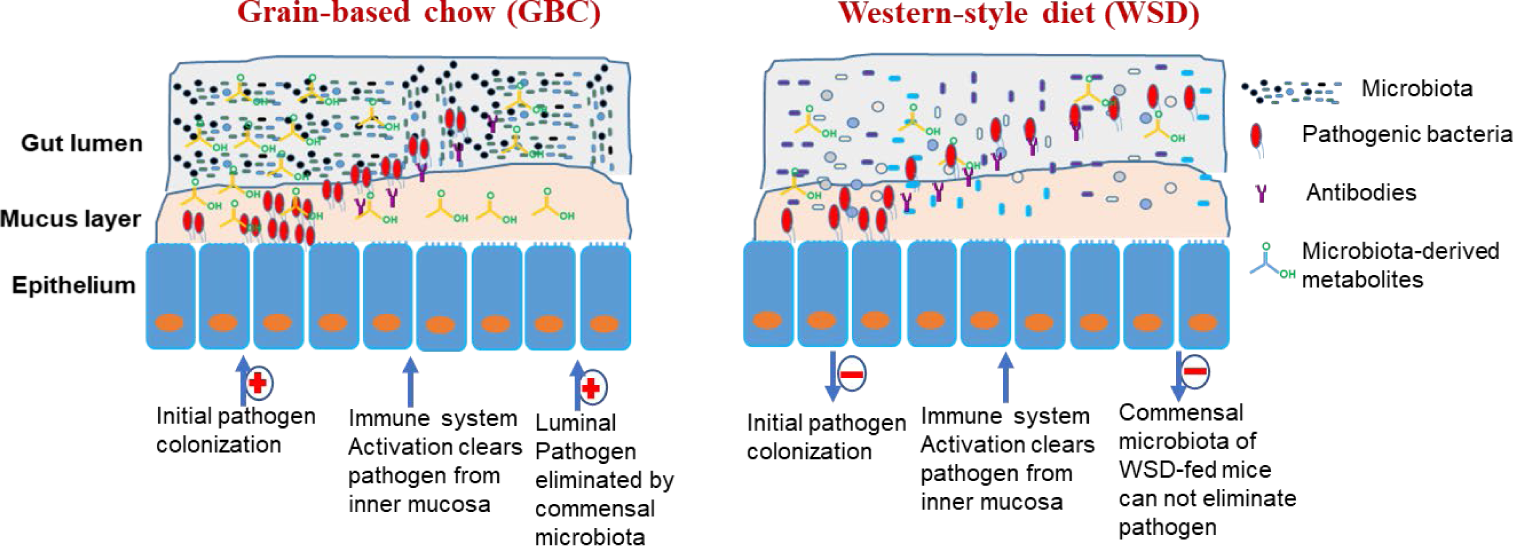

## INTRODUCTION

The intestinal tract is densely colonized by a diverse group of microorganisms collectively referred to as gut microbiota [1]. A major benefit of gut microbiota is to protect its host from infection by bacterial pathogens [2]. Such protection is mediated by ability of microbiota to directly impede colonization of pathogens via niche occupation, production of bacteriocins, and consumption of essential nutrients, i.e. colonization resistance [3]. Additionally, microbiota drives development, and regulates tone of, the mucosal immune system, which is critical to survival in response to challenge by many bacterial pathogens [4-6]. The importance of microbiota in defense against gut pathogens is readily appreciated by the stark increase in proneness to a variety of pathogens, including *Salmonella* and *C. difficile*, following antibiotic treatment [7, 8]. Use of a tractable model to study the role of the microbiota, namely use of germfree mice, also indicate a critical role of the microbiota in mediating clearance of gut pathogens from the intestinal lumen while host immunity cleared the pathogen from the epithelium [9, 10].

It is increasingly appreciated that, irrespective of antibiotics, gut microbiota composition is influenced by a variety of environmental (i.e. non-genetic) factors, especially diet [11, 12]. The majority of microbiota reside in the cecum and colon thus relying upon digestion-resistant complex carbohydrates, i.e. fiber, as a carbon source [13]. Furthermore, a variety of other dietary components are not completely absorbed and thus present in variable amounts that can modulate microbiota composition and function. Consequently, alteration of diet, can impact absolute numbers of gut bacteria (i.e. microbiota density) and influence its species composition, gene expression and localization [14-18]. Indeed, it has been proposed that broad changes in societal dietary habits, especially consumption of processed foods that are frequently rich in fats and simple carbohydrates but lacking in fiber may have impacted microbiota in a manner that promotes a variety of chronic inflammatory diseases such as inflammatory bowel disease, metabolic syndrome, and cancer [19-22]. While the nature of such changes are complex and multifactorial, we hypothesize that a well nourished microbiota drives enterocyte proliferation, mucus secretion, and IL-22-mediated production of antimicrobial peptides, which collectively maintain a robust barrier that prevents microbiota encroachment, thereby avoiding inflammation [17]. Accordingly, we postulate that reduced nourishment of microbiota, particularly due to lack of dietary fiber, has contributed to the post mid-20^th^ century increased prevalence of chronic inflammatory diseases such as inflammatory bowel disease, metabolic syndrome, and cancer.

Ability of diet to impact microbiota and, consequently, host immunity can also be envisaged to impact outcomes following encounters with gut pathogens. For example, the reduction in microbiota density resulting from a low-fiber diet might reduce colonization resistance and impede host defense, including reduced production of mucus and antimicrobial peptides. Furthermore, lack of fiber results in microbiota digesting mucus further weakening its ability to impede pathogens [13]. On the other hand, starving microbiota via lack of fiber, might limit availability of nutrients required by pathogens thus impeding pathogens colonization. Additionally, encroachment of microbiota to mucosa might limit access of pathogens, to the epithelial cells they colonize. Moreover, altered fiber content is but one change in dietary habits that might impact host-pathogen interactions. Indeed, reduced consumption of dietary fiber is associated with increased consumption of fat and simple carbohydrates. Hence, the central goal of this study was to investigate how change from a relatively unrefined grain-based chow (GBC) diet to a highly processed high-fat western style diet (WSD) impacted infection with 2 well-studied gut pathogens. We observed that the colonic environment in WSD-fed mice impeded initial colonization by pathogens but also failed to support their microbiota-mediated clearance.

## RESULTS

### WSD feeding altered gut microbiota composition and dynamics of Citrobacter infection

In accord with previous studies [17], switching mouse diet from standard grain-based rodent chow (GBC) to a high-fat low-fiber compositionally-defined western-style diet (WSD) resulted in a rapid stark (≈10-fold) reduction in the total number of bacteria per mg feces (Fig 1A), which associated with altered gut microbiota composition as indicated by PCoA (Figure 1B) and taxonomic analysis (Figure 1C), including increased alpha diversity as measured by Faith’s phylogenic diversity (Figure 1D). To investigate if such changes impacted proneness to a gut pathogen, mice were orally inoculated with *Citrobacter rodentium*. Relative to GBC-fed mice, those fed WSD initially displayed lower levels of fecal *C. rodentium* particularly at early time-points (Fig 1E). Yet, the most striking consequence of WSD consumption was frequent inability to clear this pathogen (Figure 1E). Specifically, *C. rodentium* was uniformly absent in feces of GBC-fed mice by 21 d post-inoculation but remained present in the majority of mice consuming WSD for the additional 5 weeks that they were monitored (Figure 1E). Furthermore, while GBC-fed mice were completely resistant to subsequent challenge with *C. rodentium*, those WSD-fed mice that had cleared *C. rodentium* remained prone to chronic infection when re-challenged by this pathogen (Fig 1F).

**Figure 1.**
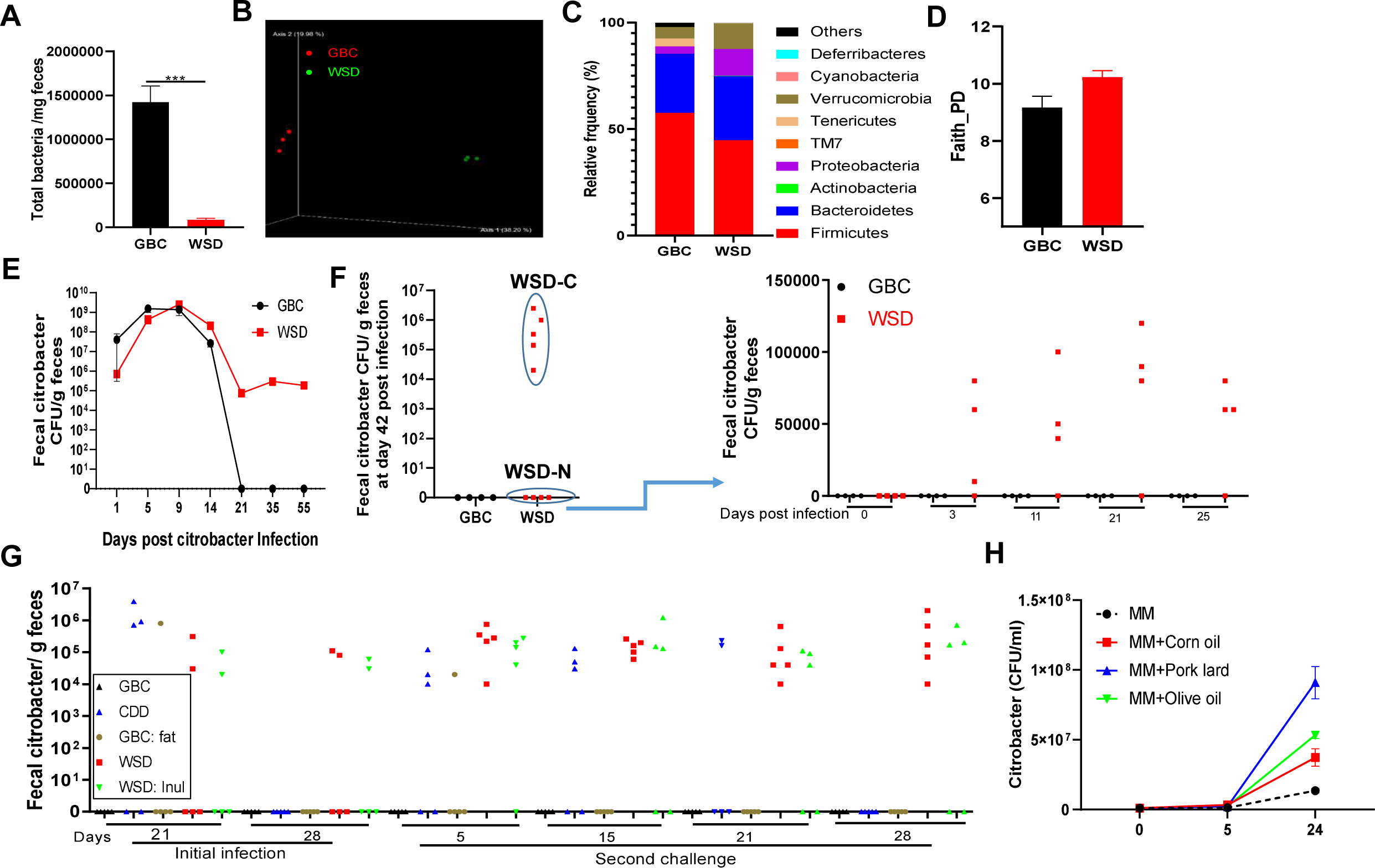
WSD-induced alteration in gut microbiota associate with inability to clear *C. rodentium*. **A-D**. Adult C57BL/6 mice were maintained on GBC or administered WSD for 1 week Bacteria per mg feces were measured by qPCR (A), gut microbiota composition was analysis by 16s rRNA sequencing: global composition as expressed by UniFrac PCoA analysis (B), relative abundance of bacterial phyla (C), Alpha-diversity (species richness) as shown by Faith PD (D). **E**. Fecal *C. rodentium* CFU was monitored. **F**. At day 42 post-inoculation, mice with and without detectable *C. rodentium* CFU, referred to respectively as WSD-C (Citro present) and WSD-N were re-inoculated with *C. rodentium*, and fecal *C. rodentium* CFU quantitated. **G**. Mice were fed indicated diet for 1 week prior to *C. rodentium* inoculation. Quantification of fecal *C. rodentium* at indicated time post initial and secondary administration of *C. rodentium*. **H**. Growth curve of *C. rodentium* in minimal culture medium supplemented with different types of lipid. Data are the means +/- SEM (n=3-10 mice per group) of an individual experiment and representative of at 2-3 separate experiments.

One primary difference between GBC and WSD is their 5% vs. 25% respective fat content (wt/wt). Thus, to specifically examine the role of such high-fat content for *C. rodentium* persistent infection, we re-proportioned the semi-purified ingredients to approximate the fat content of GBC. Consumption of this low-fat WSD recapitulated delayed clearance of *C. rodentium* and proneness to re-infection but yet such mice eventually cleared *C. rodentium*. Moreover, enriching GBC with the same lipid used in WSD formulation partially mimicked the WSD phenotype (Figure 1G). *In vitro, C. rodentium* demonstrated ability to utilize lipid as a sole carbon source in that polyunsaturated fat (corn oil), monounsaturated fat (olive oil) and animal fat (pork lard) all fueled its growth in minimal media (Fig 1H). Together, these results indicate that the high-fat content contributes to, but it not sufficient to fully recapitulate *C. rodentium* persistence. Another key feature of WSD is its low fiber content (5% by wt) and complete lack of fermentable fiber whereas GBC contains about 20% fiber, including similar amounts of fermentable and non-fermentable (insoluble) fiber. Enriching WSD with the fermentable fiber inulin can restore many aspects of host-microbiota homeostasis, including bacterial density, and prevent WSD-induced metabolic syndrome [17, 23]. Enriching WSD with inulin did not significantly aid clearance of *C. rodentium* (Figure 1G). Thus, inability of WSD-fed mice to manage *C. rodentium* can not be easily corrected by simply increasing bacterial density with inulin and, rather, likely reflects impacts of multiple ingredients, and/or lack thereof, in this diet.

### Low virulence of *C. rodentium* in WSD mice

Next, we examined the extent to which WSD influenced *C. rodentium*-induced disease. Mice consuming GBC or WSD were euthanized 10d following *C. rodentium* inoculation, at which time levels of *C. rodentium* were similar on the 2 diets, or 42d, at which time only WSD mice remained infected with *C. rodentium*. Levels of inflammatory marker fecal lipocalin-2 peaked at 10d in both groups and were moderately lower at all times in WSD-fed mice (Figure 2A). Accordingly, WSD-fed mice exhibited reduced indications of acute inflammation as reflected by splenomegaly (Figure 2B), colon shortening (Figure 2C), colonic hyperplasia (crypt lengthening) (Figure 2D) and depletion of goblet cells (Figure S1), all of which are typical features of *C. rodentium* infection. Moreover, the moderate degree of gut inflammation observed in WSD-fed mice resolved with similar kinetics as that of GBC-fed mice. WSD-fed mice that had or had not cleared *C. rodentium* at 42 d post infection, lacked abnormal mucosal pathology such as goblet cell depletion, which is often observed in chronic infection (Figure 2&S1). Such lack of chronic overt inflammation in WSD-fed mice suggested lack of virulence gene expression by persisting C. rodentium. Indeed, mucosal-adherent *C. rodentium* was present at 10d, but not 26d following its inoculation in mice fed either diet, despite its persistence in feces of WSD-fed mice (Figure 3A&B). Accordingly, expression of two *C. rodentium* key virulence genes, namely Ler and Tir, normalized for fecal levels of *C. rodentium*, was markedly reduced in WSD-fed mice at the later time point (Figure 3C&D). Moreover, fecal transplant from the persistently-infected mice to GBC-fed immune deficient mice (Figure 3E), resulted in only modest levels of C. rodentium in the new host further supporting the notion that the persisting *C. rodentium* was not highly virulent (Figure 3F&G).

**Figure 2.**
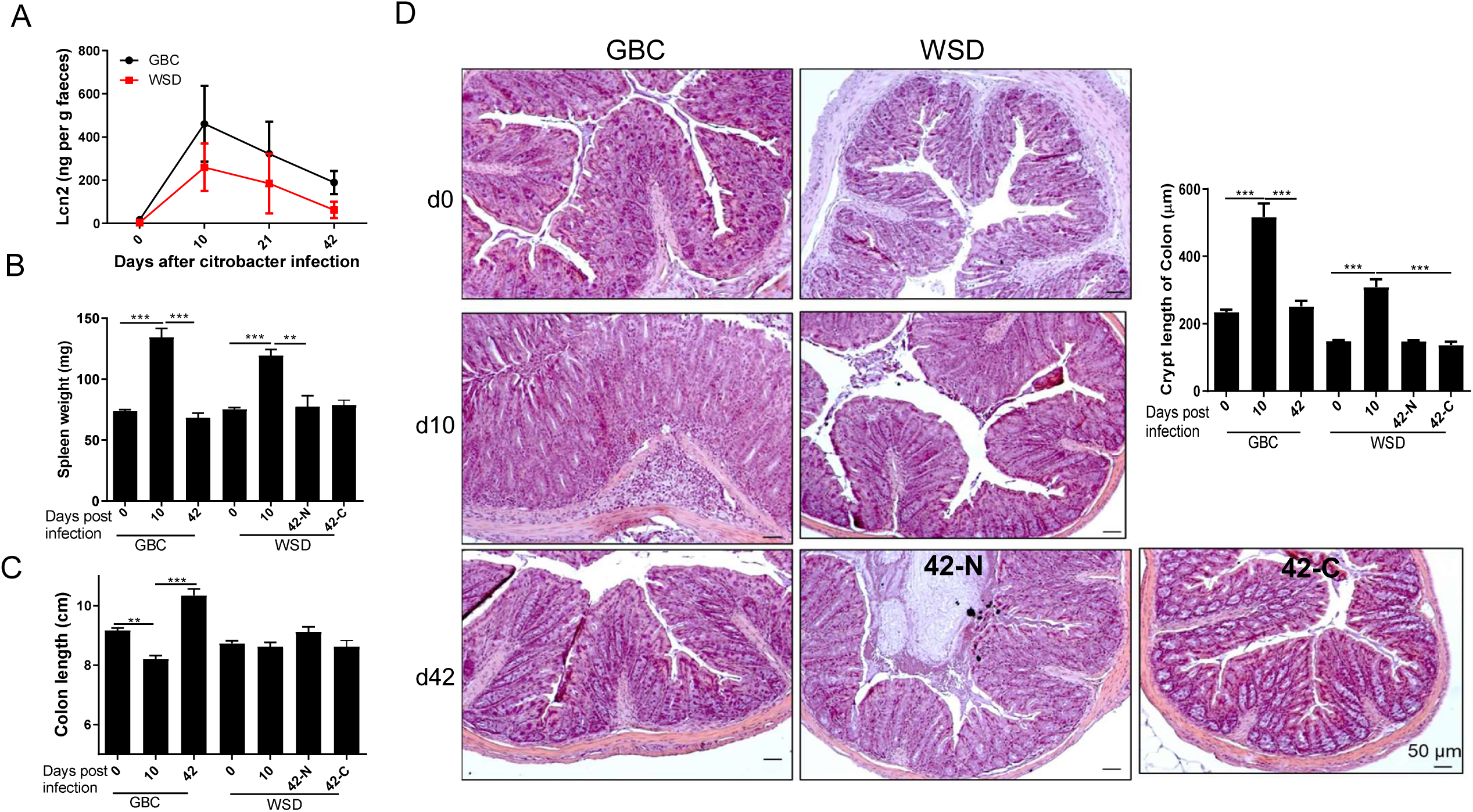
Reduced acute inflammation and lack of overt chronic inflammation in WSD-fed mice challenged with *C. rodentium*. C57BL/6 mice (n=3-10 per group) were administered PBS *C. rodentium* and euthanized at indicated time points. **A**. Measurement of fecal lipocalin 2 by ELISA. **B-C**. Spleen weight (B) and colon length (C) was measured in GBC and WSD fed mice that had (WSD-C) or lacked (WSD-N) detectable fecal *C. rodentium* CFU. **D-E**. Colon tissue was processed for H&E staining, which afforded measure of colon crypt length.

**Figure 3.**
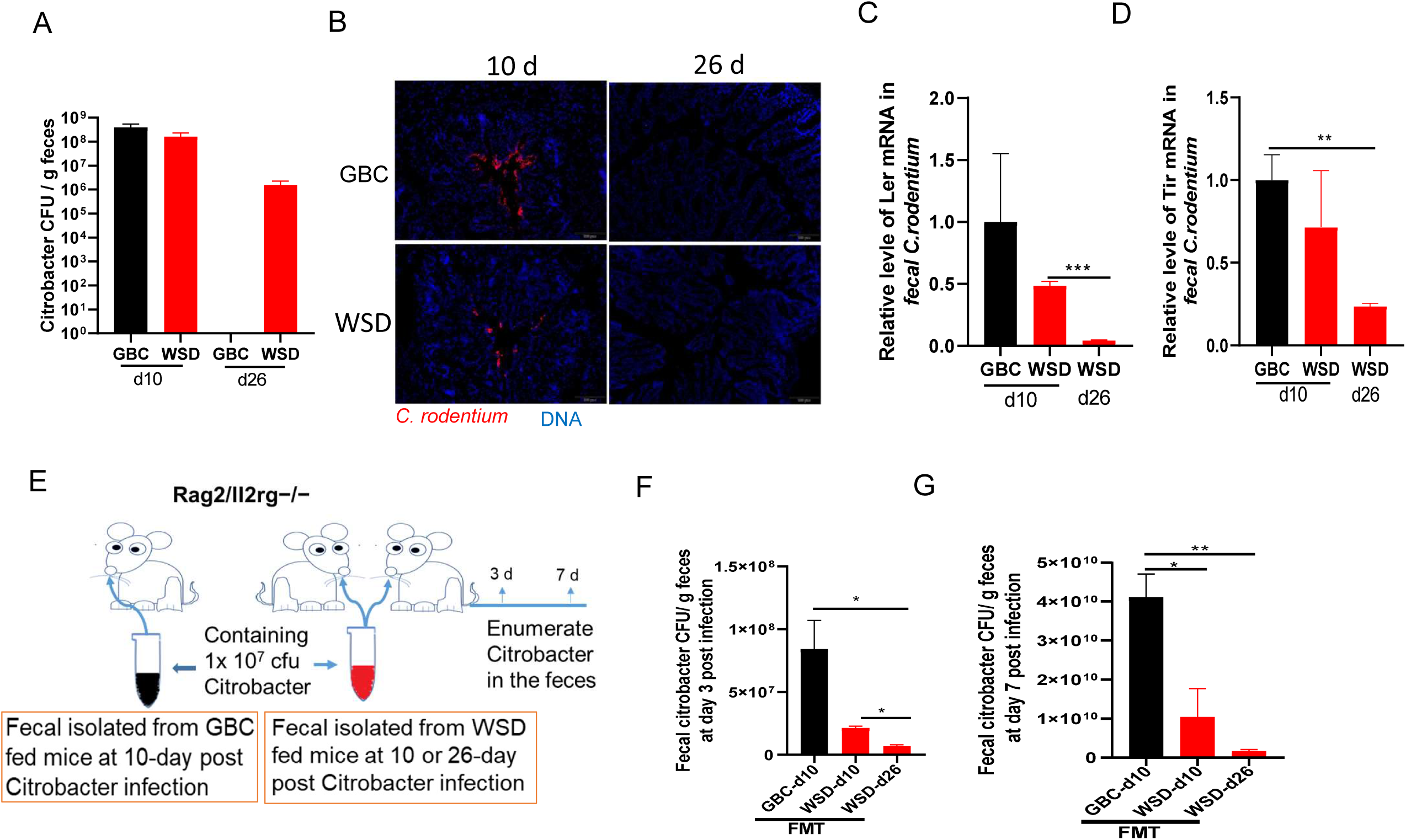
WSD feeding resulted in reduced virulence gene expression by C. rodentium. **A**. Quantification of fecal *C. rodentium* at indicated days post-inoculation. **B**. Immunofluorescent staining of *C. rodentium* in colon tissue at indicated days post-inoculation. **C-D**. The relative expression level of *C. rodentium* virulence genes measured by qRT-PCR. **E-G**. Rag1^-/-^IL2R^-/-^ mice were orally administered fecal suspension containing 1×10^7^ CFU *C. rodentium* as schematized (E). Quantification of fecal *C. rodentium* at day 3 (F) and 7 (G) post inoculation. Data are the means +/- SEM (n=3-5 mice per group)

### Persistent *C. rodentium* infection promotes insulin resistance

It is increasingly appreciated that some host-microbial interactions that don’t cause overt, i.e. histopathologically-evident, inflammation can promote pro-inflammatory gene expression that promotes insulin resistance, especially in the context of WSD-induced obesity [24]. Hence, we compared gene expression and metabolic parameters in WSD-fed mice that had or had not cleared *C. rodentium*. At 42 d post *C. rodentium* infection, *C. rodentium* persistence associated with increased expression of inflammation-related genes in colon (Figure 4A-D) but not insulin resistance (Figure 4E). However, glycemic assessment of these mice 5 months post-inoculation found increased fasting blood glucose levels and reduced responsiveness to insulin in the mice that failed to clear *C. rodentium* (Figure 4F-H) despite similar weight and adiposity (Figure S2A-E). Such insulin resistance was associated with elevated expression of canonical markers of low-grade inflammation, namely IL-6 and TNFα, in adipose tissue and liver (Figure 4I-L)

**Figure 4.**
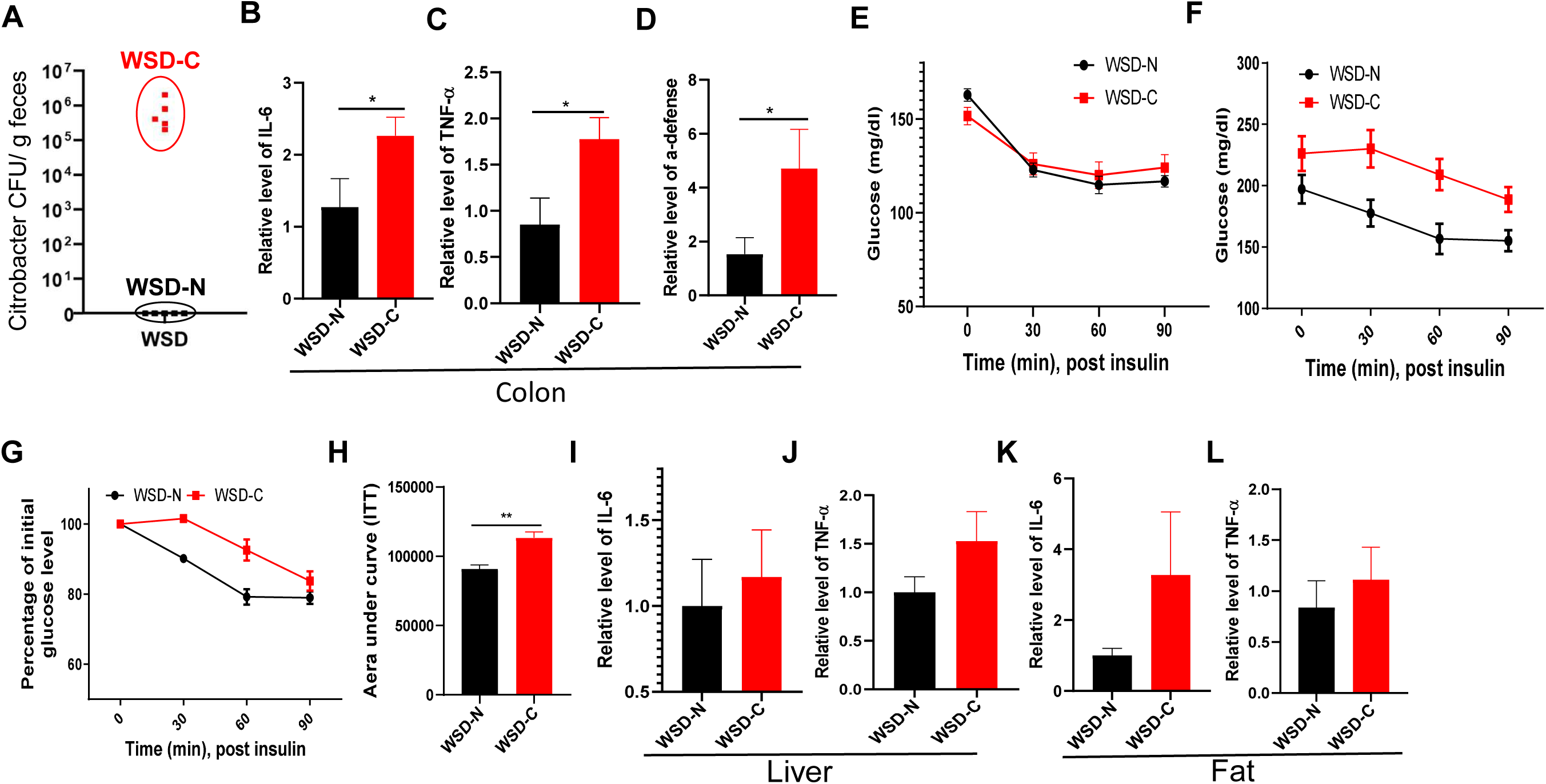
Persistence of Citrobacter infection in WSD-fed mice associates with low-grade inflammation and insulin resistance. **A**. Measure of fecal *C. rodentium* CFU was used to stratify WSD-fed as being persistently infected with (WSD-C), or having cleared (WSD-N), *C. rodentium* 42 days-post-inoculation, at which time we measured colonic expression of IL-6 (B), TNF-α (C) and α-defensin (D), by qRT-PCR, and glycemic responses to insulin. **F-K**. 5 months later, glycemic responses and indices of low-grade systemic inflammation were measured. Measure of absolute (F) and relative (G) glucose levels following insulin as well as area under curve were calculated (H). Relative level IL-6, TNF-α in liver and epididymal fat were analyzed by qPT-PCR (I-L). Data in B-L are means +/- SEM of an experiment with 10 mice that, as shown in A, yielded an n=5 for B-L.

The association of chronic *C. rodentium* infection and dysglycemia suggested a potential role for persistent *C. rodentium* infection in dysglycemia and/or that dysglycemia impeded clearance of *C. rodentium*. To help differentiate these non-mutually exclusive possibilities, we examined if streptozotocin-induced diabetes impacted the course of *C. rodentium* infection. We selected mice in which STZ elevated fasting glucose to levels only slightly higher than those of *C. rodentium*-infected WSD-fed mice (Figure S3A). Such hyperglycemia associated with delayed but nonetheless eventually complete clearance of the pathogen (Figure S3B). These results suggest dysglycemia contributes to, but does not fully account for, the impaired clearance of *C. rodentium* by WSD-fed mice. We next examined the extent to persistent *C. rodentium* in WSD-fed mice might reflect impaired mucosal immunity. WSD, by itself, results in lower expression of IL-22, which is known to play an important role in protection against *C. rodentium*. However, IL-22 expression was higher in mice in persistently infected WSD-fed mice than in GBC-fed mice that had fully cleared this pathogen (Figure S4A). Analogously, WSD-fed mice displayed elevated level of fecal and serum antibodies to *C. rodentium* arguing against the notion that defective adaptive immunity under *C. rodentium* accounted for the persistent infection (Figure S4B&C).

### WSD-induced changes in microbiota contribute to *C. rodentium* persistence

While adaptive immunity is required to clear epithelial-adherent *C. rodentium*, elimination of this pathogen from the outer mucus and lumen is thought to result from *C. rodentium* being out-competed by commensal microbiota [9]. Hence, we hypothesized that inability of WSD-fed mice to eliminate *C. rodentium*, might reflect low microbiota density, altered species composition, and/or lack of inflammation-induced changes in microbiota, which may help clear oxygen-using pathogens [25]. To test this hypothesis, we measured *C. rodentium*-induced changes in microbiota infection via qPCR and 16s rRNA sequencing (Figure 5 and Figure S5). The total bacteria per mg feces increased slightly in GBC-fed mice while modestly decreasing in WSD-fed mice following administration of *C. rodentium* (Figure 5A). Moreover, PCoA analysis of 16S sequencing data revealed that extent and duration of *C. rodentium*-induced changes microbiota composition was also quite different between mice fed GBC and WSD. Specifically, while GBC-fed mice displayed a stark change in microbiota composition at 42 post-inoculation, at which time they had uniformly cleared the pathogen, WSD fed mice showed only modest changes at this time regardless of whether they had cleared the pathogen (Figure 5B&C). Such relatively reduced changes in microbiota in response to *C. rodentium* in WSD-fed mice were also evident by use of the Linear discriminant analysis effect size (LefSe) algorithm (Figure S5D-F). Moreover, assessment of alpha-diversity (i.e. species richness) by Faith’s phylogenetic diversity and number of observed taxonomic units (OTU) showed increased community richness, which was not observed in WSD-fed mice (Figure 5D-F). While analysis at phylum and major genera levels analysis revealed basal differences in mice fed WSD and GBC, differences the relative changes in response to *C. rodentium* were similar at this level although, among WSD-fed mice, failure to clear the pathogen was associated with reduced *Proteobacteria*, in part driven by reduced levels of *Desulfovibrio* (Fig S5 A-C).

**Figure 5.**
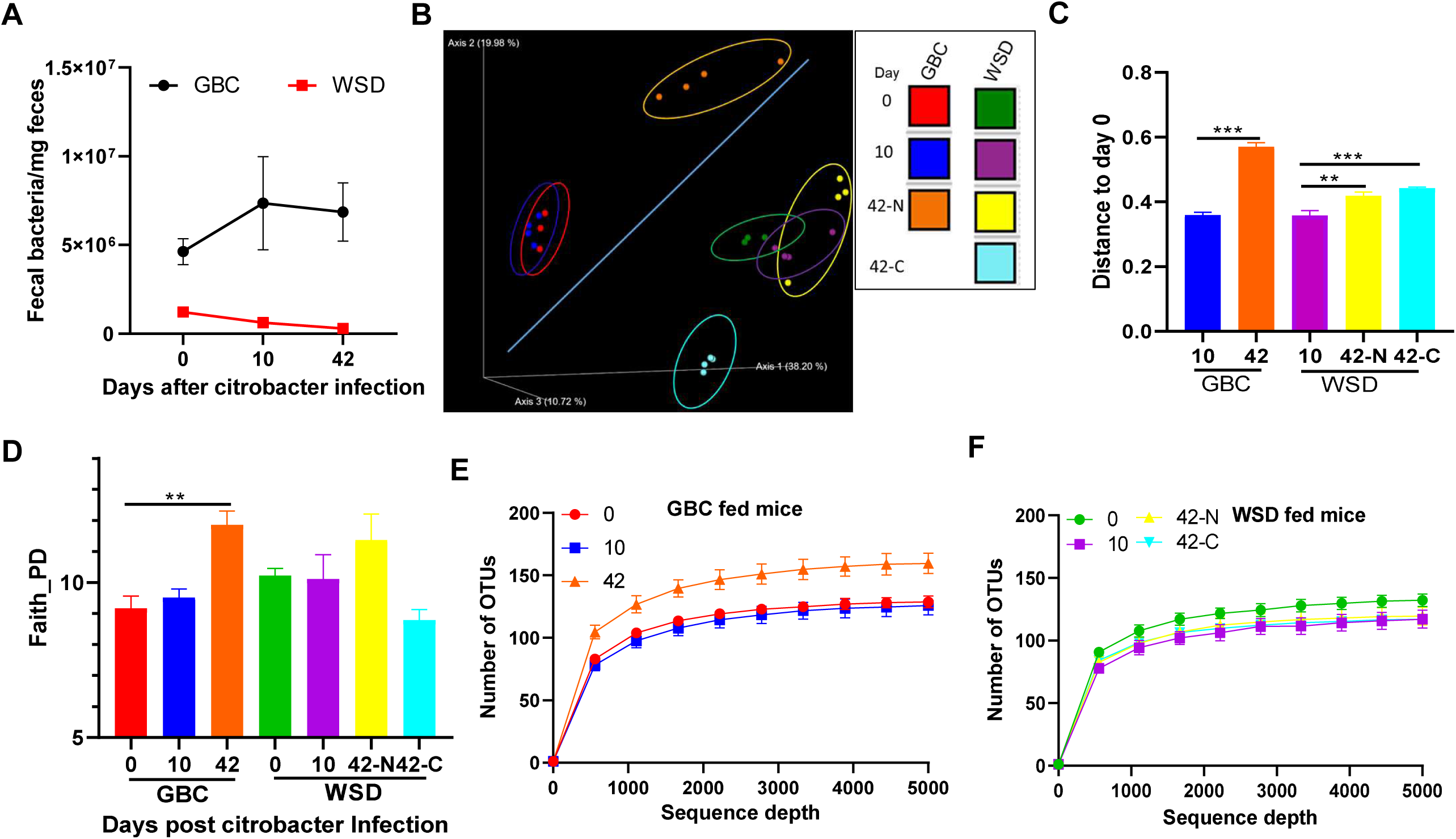
*C. rodentium* starkly alters microbiota of mice fed GBC but not WSD. **A**. Measure of total fecal bacterial DNA by qPCR relative to time of *C. rodentium* administration. **B-F**. Analysis of gut microbiota composition based on 16S RNA sequencing and analyzed/displayed by unweighted UniFrac PCoA (B), UniFrac distances between Day 10 or 42 after *C. rodentium* relative to Day 0 (C). Alpha-diversity as measured by Faith PD (D) and the observed OTUs (E&F).

To investigate the extent to which differences in microbiota in mice fed GBC and WSD might contribute to *C. rodentium* persistence, we utilized gnotobiotic mice with a very limited microbiota, specifically the eight bacterial species known as the Altered Schaedler flora (ASF). Unlike germfree mice, such ASF mice have relatively normal immune and metabolic functioning [26]. We observed that, irrespective of whether they were fed GBC or WSD, ASF mice were unable to clear *C. rodentium* infection as evidenced by uniformly high levels of the pathogen in their feces 5 weeks post-inoculation (Figure 6A). Administration of feces from conventional mice, which were maintained on GBC, to such chronically infected mice resulted in complete clearance of the pathogen in the recipients fed GBC but resulted in only a moderate and transient lowering of *C. rodentium* CFU in those fed WSD (Figure 6B). This result suggests that a properly nourished complex microbiota promotes clearance of *C. rodentium* from the intestinal tract. To test this possibility we sought to mimic the WSD-induced reduction in gut bacterial density via a 1-week course of kanamycin in GBC-fed mice, which we administered 15 days following inoculation with *C. rodentium*, at which time rapid clearance of fecal CFU would typically commence. This kanamycin regimen delayed clearance of *C. rodentium* until 2-3 weeks after cessation of the antibiotic (Figure 6B). Furthermore, administering kanamycin to GBC-fed mice following initial clearance of *C. rodentium* infection made those mice prone to re-infection by this pathogen, which was only cleared after cessation of the antibiotic (Figure 6C). Analogous results were obtained when these approaches were used to investigate the susceptibility to a second challenge with *C. rodentium*, Specifically, while mice maintained continuously on GBC were impervious to re-infection when re-challenged by *C. rodentium* 2 weeks following initial clearance of this pathogen, those switched to WSD 7d following its initial clearance were prone to re-infection. Levels of *C. rodentium* in these mice were markedly augmented by kanamycin administered 3 weeks post-inoculation of *C. rodentium* (Figure 6D). Together, these results argue that persistence of *C. rodentium*, and proneness to re-infection, in WSD-fed mice likely results from their alterations in gut microbiota, particularly their low bacterial density.

**Figure 6.**
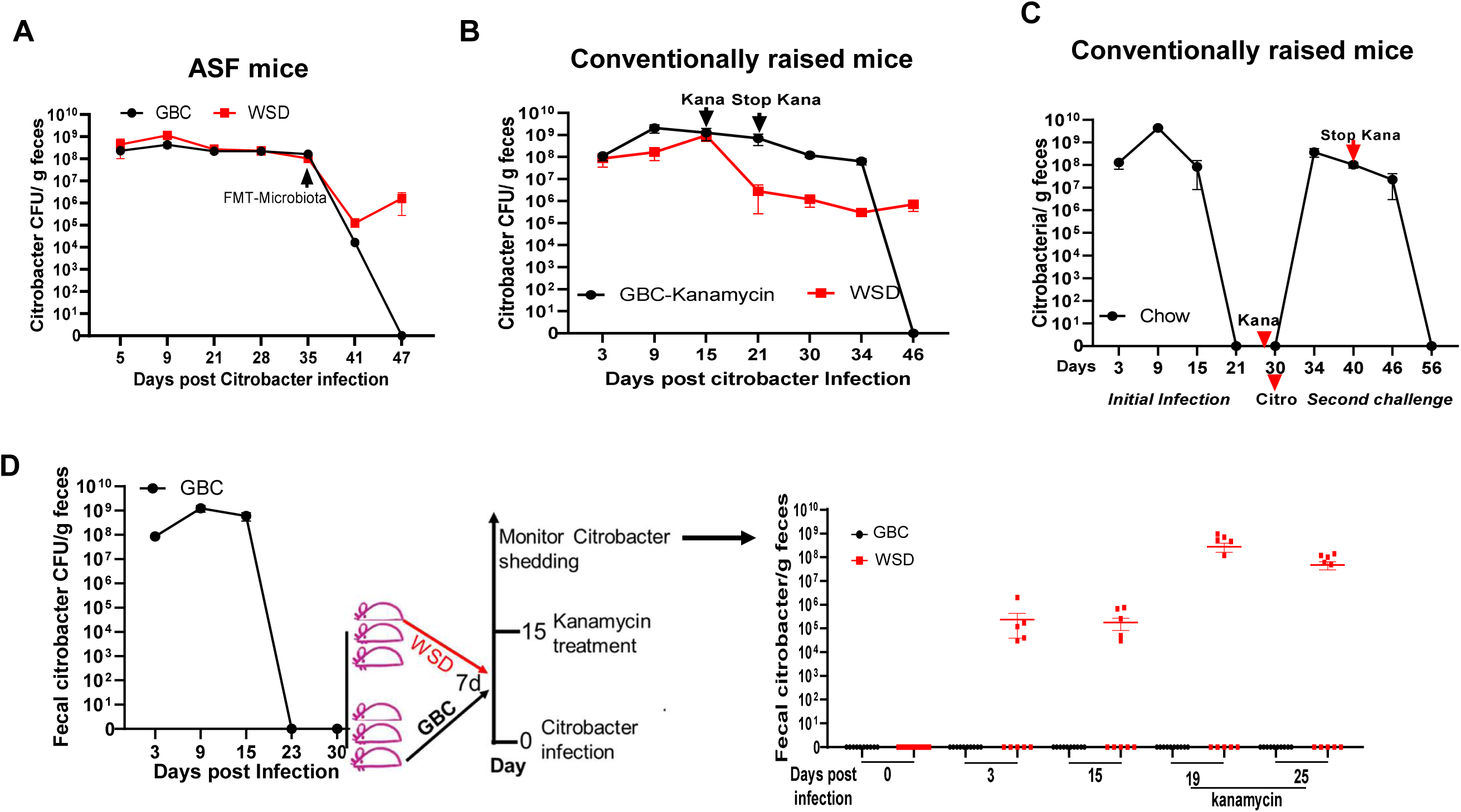
A complex microbiota, nourished by GBC mediates clearance of *C. rodentium* persistent infection. **A**. ASF mice were fed GBC or WSD for 1 week before orally inoculation with *C. rodentium*. 35d later, these mice were administered fecal suspension prepared from conventional GBC-fed mice. **B**. GBC-fed conventionally-raised C57BL/6 mice were orally administered *C. rodentium*. 15 days later, mice were left untreated or orally administered kanamycin (1000mg/kg) on days 15, 17, 19, and 21. **C**. GBC-fed, *C. rodentium* infected mice were orally garaged with kanamycin at day 29 post-inoculation, and re-administered *C. rodentium* at day 30, kanamycin was administered every other day from day 21-29. **D**. GBC-fed mice were administered *C. rodentium*, which they uniformly cleared by 30d, at which time they were maintained on GBC or switched to WSD. One week later, mice were re-challenged with *C. rodentium* and treated with kanamycin 15 d later. All data shown is measure of fecal C. rodentium CFU on indicated day in that experiment. Data are the means +/- SEM (n=3-10 mice per group)

### WSD supplemented with fermentable fiber inulin promote *C. rodentium* initial colonization

We next turned our attention to early events in *C. rodentium* infection, specifically investigating why, despite their low-microbiota density, which one would presume should lower colonization resistance, WSD-fed mice exhibited reduced colonization and inflammation shortly following initial exposure to the pathogen. We first probed a role for mucosal immunity, in particular, considering that WSD impacts IL22, IL-18, and adaptive immunity, all of which impede *C. rodentium* infection. However, WSD’s reduction of initial C. rodentium colonization was not impacted by genetic deficiency of IL-22, IL-18, or adaptive immunity (Rag1^-/-^) thus arguing against a role for these host factors (Figure S6). Hence, we considered the possibility that WSD’s lack of fermentable fiber might have resulted in a nutrient-poor colon environment that impeded C. rodentium growth. In accord with this possibility, we observed that enrichment of WSD with fermentable fiber, inulin, resulted in more uniform colonization in response to low inoculums of *C. rodentium* (Figure 7A) and, moreover, increased levels of *C. rodentium* to those of GBC-fed mice (Figure 7B). Such restoration of fecal CFU was associated with more abundant staining for *C. rodentium* in colonic mucosa (Figure 7C) as well as greater indices of inflammation, including splenomegaly (Figure 7D) and levels of fecal Lcn-2 (Figure 7E). The notion that inulin directly impacted the ability of the colon to support C. rodentium growth was tested in vitro. Directly supplementing minimal media with inulin did not promote *C. rodentium* growth indicating that *C. rodentium* is not able to directly utilize inulin as a carbon source (Figure 7F&G). However, enriching minimal media with fecal and cecal supernatants from mice fed inulin-enriched WSD enhanced growth of *C. rodentium* (Figure 7H). Dietary inulin is known to be metabolized into short-chain fatty acids, which we tested for ability to fuel *C. rodentium* growth in vitro. We observed acetate, but not butyrate or propionate was able to support *C. rodentium* growth (Figure 7I). Together, these results suggest that fermentation of fiber by commensal bacteria can fuel rapid colonization of C. rodentium following its oral ingestion.

**Figure 7.**
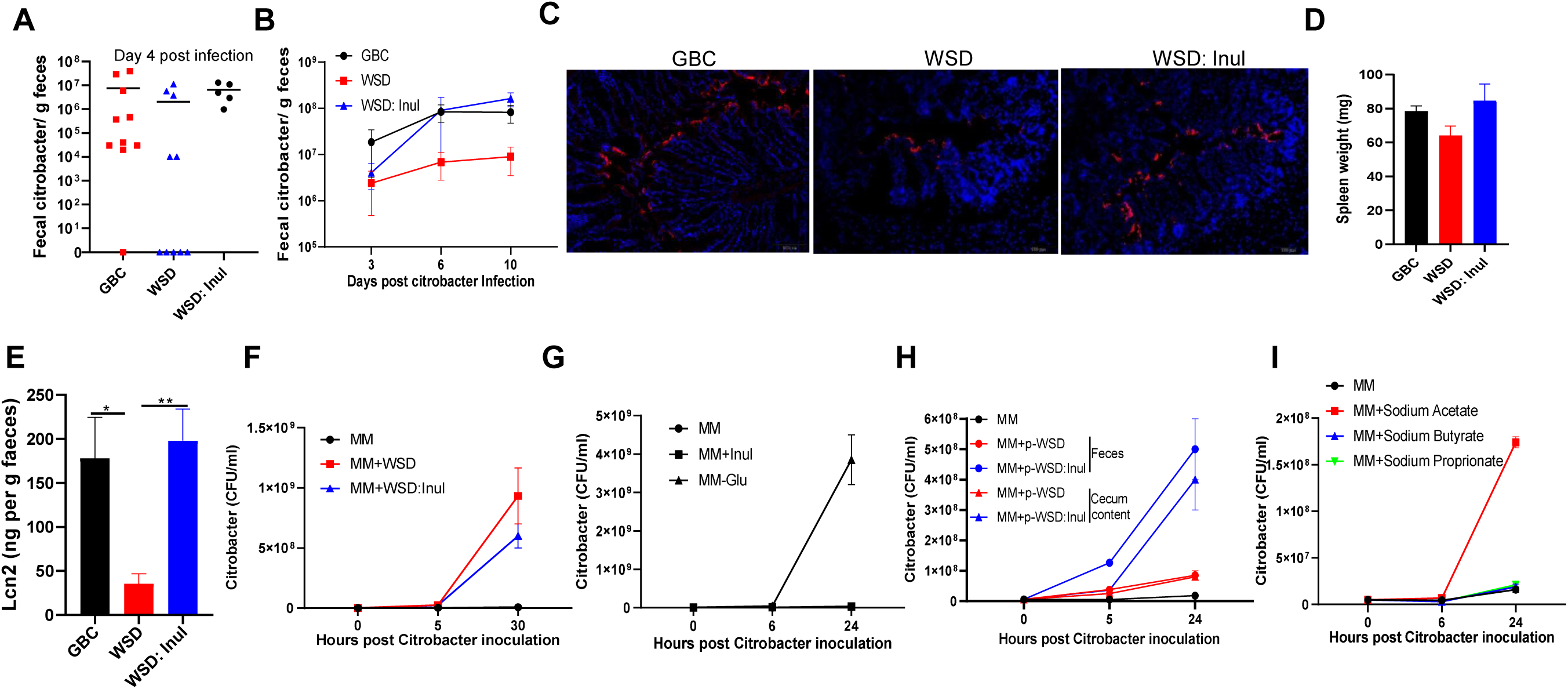
Fermentable dietary fiber promotes initial colonization of *C. rodentium*. **A**. Mice fed with indicated diets (n=5-10) were orally administered 2×10^8^ CFU *C. rodentium* per mouse and fecal *C. rodentium* CFU measured 4 days later. **B**. Mice were administered with a dose of 4×10^*8*^ CFU *C. rodentium /mouse*, fecal *C. rodentium* CFU measured at indicated times. **C-E**. Mice were euthanized on day 10 post-inoculation. Immunofluorescent staining of *C. rodentium* in colon (C), Spleen weight (D), Fecal Lcn-2 (E). **F-I**. In vitro growth of *C. rodentium* in minimal medium supplemented with WSD diets +/- inulin (F), pure inulin (Inul) or glucose (Glu) (G), fecal or cecal supernatant from mice fed WSD +/- inulin (H), or indicated purified short-chain fatty acid.

### WSD supplemented with fermentable fiber inulin promote *Clostridium difficile* initial colonization but improved the clinical-type consequences

Lastly, we examined the extent to which our findings using *C. rodentium* might be applicable to another colitis-inducing pathogen, namely *Clostridium difficile*, to which mice are made prone to by a cocktail of antibiotics [27]. We observed that, relative to GBC-fed mice, WSD feeding delayed initial colonization of *C. difficile*, which was associated with a delay and reduction in clinical-type consequences, namely death and weight loss (Figure 8A-C). Enriching WSD with inulin largely, but not entirely, restored rapid *C. difficile* colonization but did not result in the rapid weight loss and uniform death seen in GBC-fed mice, nor the 60% mortality exhibited by WSD-fed mice (Figure 8A-C). Levels of *C. difficile* CFU and toxin were well correlated and matched the kinetics of inflammation/tissue damage displayed by mice fed each diet (Figure 8D-F). Measure of total gut bacteria by qPCR indicated that despite the large amounts of antibiotics used in this model, WSD still reduced initial bacterial density, which was largely restored by inulin enrichment (Figure S7A). WSD also impacted microbiota composition in this context, which was not restored by inulin (Figure S7B-D). Together, these results suggest a high potential for fermentable dietary fiber to promote growth of *C. difficile* in the colon and perhaps influence microbiota composition and host responses in a manner that influences outcomes of exposures to this pathogen.

**Figure 8.**
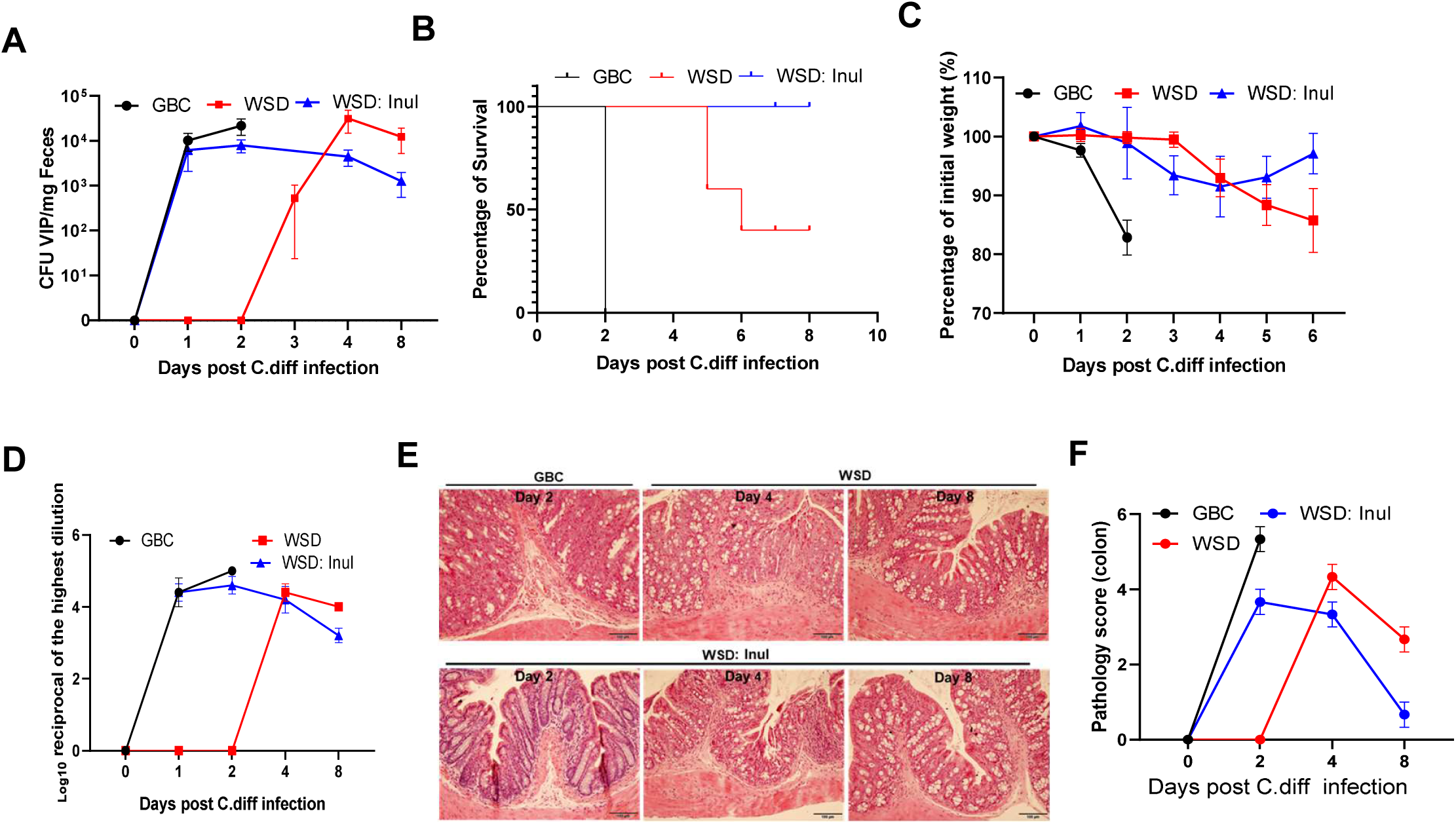
WSD impedes initial *C. difficile* colonization. **A**. Quantitative PCR was used to quantify the *C. difficile* loads in antibiotic-pretreated wild type mice (n=3-11 per group). **B**. Survival curve of antibiotic-pretreated mice fed with different diets during *C. difficile* infection. **C**. Body weight measured and percentage to initial weight calculated. **D**. Cytotoxin in feces was measured by Vero cell cytotoxic assay. **E&F**. Histological analysis of proximal colons. Mice were euthanized on indicated day. Representative H&E staining (E) and pathological scores (F).

**Figure 9.** Graphic summary and model. The low-fiber content of WSD impedes initial colonization of pathogens. Immune system functions appropriately irrespective of diet to clear pathogens from the mucosa. Commensal microbiota of mice fed GBC but not WSD is sufficient to eliminate pathogen from the lumen of the intestine. Underlying mechanisms discussed in text.

## DISCUSSION

The importance of the commensal microbiota in protecting against pathogenic bacteria is underscored by the markedly elevated proneness to infection that results from broad-spectrum antibiotics, which typically reduce total bacterial loads by orders of magnitude and dramatically alter its phylogenic composition [28]. Herein, we report that outcomes of exposure to bacterial pathogens are also impacted by the more moderate changes in microbiota that can result from changes in diet. Specifically, we observed that relative to grain-based chow (GBC), which is typically consumed by lab mice, feeding “western-style” diet (WSD) reduced initial colonization of 2 distinct pathogens, namely attaching-effacing enteropathogenic *E. coli*-like gram negative *C. rodentium* and gram-positive human pathogenic isolate of *C. difficile*. However, WSD-feeding also impaired clearance of these pathogens resulting prolonged infections. Such complex altering the dynamics of these infections suggests that diet may be a key determinant of the heterogenous course/outcomes of such infections and/or the evolving epidemiology of these infectious diseases.

The reduced ability of pathogens to rapidly colonize the colon of WSD-fed likely reflects, at least in part, this diet’s lack of fermentable fiber in that enriching WSD with the fermentable inulin fully restored rapid colonization by *C. rodentium* and *C. difficile*. We envisage 2 potential non-mutually exclusive mechanisms whereby lack of fermentable fiber might impede pathogen initial colonization. Most simply, we postulate that lack of fiber reduces nutrients available to colon bacteria thus heightening competition for these nutrients. In support of this possibility, we note that commensal Bacteroides, namely B. thetaiotamicron, will outcompete C. rodentium when only simple carbohydrates are available but will promote C. rodentium growth in mice fed GBC [9]. In this scenario, enriching WSD with inulin will ease competition for the simpler carbohydrates required by C. rodentium and, in that course, produce metabolites such as SCFA that can be directly utilized by the pathogen. Accordingly, these results suggest that the delayed increase in both commensal bacterial density and *C. difficile* levels in mice consuming inulin-enriched WSD reflects the large depletion of commensal bacteria by the antibiotic cocktails, utilized in the *C. difficile* but not *C. rodentium* infection models, delayed metabolism of inulin by those bacteria that withstood antibiotics. Another possibility is that the microbiota encroachment that results from low-fiber diets including WSD [17], creates a localized colonization resistance within the inner mucus, which pathogens such as C. rodentium must traverse to adhere to epithelial cells.

Mobilization of the immune system, especially adaptive immunity, efficiently clears many pathogens, including C. rodentium from the epithelium and inner mucus thus making its persistence fully dependent upon outcompeting commensal bacteria for nutrients. Accordingly, we found that inability of WSD-fed mice to clear *C. rodentium* was not associated immunity but, rather, largely reflected inability to clear this pathogen from the intestinal lumen. Thus, the stark changes in microbiota density and composition induced by WSD are likely germane the persistence of *C. rodentium* in WSD-fed mice. That a low-fat version of WSD, generated by reproportioning the semi-purified components of WSD to match the approximate macronutrient composition of GBC partially rescued impaired clearance of *C. rodentium* suggests that the lipid component of WSD, lard, contributes to this phenotype. However, enriching GBC with lard did not have a marked impact on clearance of *C. rodentium* indicating this aspect of WSD was not the predominant driver of this phenotype. Enriching WSD with inulin did not correct the impaired clearance of, nor proneness to re-infection by, *C. rodentium* exhibited by WSD-fed mice indicating that these aspects of the WSD-induced phenotype could not be explained purely by the reduction in total bacterial density resulting from this diet. Considering that inulin’s enrichment of only microbiota density, not diversity observed in our previous study, which may be due to that inulin can only promote the growth of a certain of bacteria species, suggested a key role for specific members of gut microbiota or overall gut microbiota including both density and diversity are needed to outcompete *C. rodentium* and thus prevent its persistence.

That ASF mice, which have a very limited microbiota but are nonetheless considered immunologically and metabolically normal [26], including being capable of microbiota-dependent catabolism of complex polysaccharides, could not clear *C. rodentium*, even when fed GBC, suggests a complex microbiota is required to eliminate *C. rodentium* from the gut lumen. Furthermore, that administering ASF a complex microbiota via FMT resulted in a rapid drop in *C. rodentium* levels irrespective of diet which ASF mice were fed but yet GBC feeding was required to sustain the decrease and achieve clearance highlights the importance of as yet undefined components of diet to support the microbiota needed to eliminate the pathogen. Indeed, the normal clearance of *C. rodentium* in GBC-fed mice is accompanied by an apparent increase in alpha-diversity (species richness), which may possibly be the result of the pathogen having created a nice for low abundance bacteria that subsequently outcompete *C rodentium*.

Analogous to results from Nunez and colleagues attained in germfree mice and antibiotic treated mice model, *C rodentium* that persists in the lumen does not express its defined virulence genes and, accordingly does not appear to elicit gut inflammation [9, 29]. Yet, as shown herein, such persisting *C. rodentium* was still associated with a phenotype, namely that of exacerbating insulin resistance induced by WSD. That the last century has witnessed a decrease in classic infectious diseases paralleled by an increase in chronic inflammatory diseases, including metabolic syndrome prompts us to speculate on possible implications of our findings. Specifically, our results suggest the possibility that reduced incidence of food-borne illness may not entirely reflect reduced presence of pathogens in industrialized food but rather that some pathogens may not find highly expansive growth conditions in hosts consuming low fibers diets. However, such pathogens, or perhaps even pathobiont like organisms such as *E. coli*, may not be readily cleared from persons consuming western-style diets and thereby promote the chronic, often low-grade, inflammation that drives a variety of diseases, including metabolic syndrome.

## Materials and methods

### Mice, Diet and *C. rodentium* infection

C57BL/6 Wild type (WT) and IL-18 KO were purchased from Jackson laboratory, and housed at Georgia State University. IL-22 KO mice, generated by Genentech, were bred and housed at Georgia State University. Rag1 KO were purchased from Jackson laboratory and housed in Helicobacter negative and SPF environment. These mice were fed with grain-based chow (GBC) (cat# 5001, LabDiet), or different compositional defined diets (CDD) (Supplemental table 1, Research Diets, Inc), and housed at Georgia State University under approved animal protocols (IACUC # A17047). Mice at 6-8 weeks old age were fed with GBC or specific CDDs for 1 week before orally gavage with 4-6 ×10^8^ CFU/per mouse *C. rodentium* strain, which are bioluminescent and kanamycin resistant. After infection, the feces were collected at indicated time points and resuspend in PBS at concentration of 100 mg/ml. After homogenization, the suspension was serial diluted, then 10 µl of 10-fold serial dilutions was plated on LB agar plate with 100 µg/mL kanamycin. After 21 days post infection, mice with clearance of *C. rodentium* was confirmed, then re-infected with *C. rodentium* with 4-6 ×10^8^ CFU/per mouse at indicated time points, or GBC fed mice were switched to fed WSD for 1 week before re-infected with *C. rodentium* at dose of 4×10^8^ CFU/per mouse. For antibiotic treatment: GBC fed mice were infected with *C. rodentium* as described above. Some of these mice were gavaged with Kanamycin (1,000 mg/kg in water) every other day from 14 days post infection until indicated time points. Other mice were orally gavaged with kanamycin at day 29 post infection, then infected with *C. rodentium* with a dose of 4-6 ×10^8^ CFU/per mouse next day. The mice were euthanized, colon length and spleen weight were measured at the end of experiment. Samples were collected and stored in formalin or -80°C until further analysis.

### *C. difficile* infection

Wild-type C57BL/6 mice at 6-week-old were fed with GBC, WSD and WSD supplemented with inulin for 1 week, before received an antibiotic cocktail including of gentamicin (0.035 mg/mL), kanamycin (0.4 mg/mL), metronidazole (0.215 mg/mL), colistin (850 U/mL) and vancomycin (0.045 mg/mL) in drink water for 3 days. After one day, these mice were intraperitoneally injected with clindamycin (10mg/kg), infected with C. difficile (1 x 10^5^ spores/per mouse) by orally gavage one day later as previous described [27]. The feces were collected, and weight loss was monitored after infection. At the end of experiment, the colon was collected for HE staining. To measure *C. difficile* burden in feces, the fecal DNA was extracted and quantitative PCRs were performed in a CFX96 apparatus (Bio-Rad) with specific primers: *C. difficile* VPI-F: TGAAGGAATAGCAAATGATCCAGA; *C. difficile* VPI-R: TTCTCTACCTTTTCCCAGCTATC according to previous described. The total *C. difficile* in the feces was calculated by using a standard curve, and expressed as CFU per mg of feces.

### Gnotobiotic mouse experiments

C57BL/6 WT germ free mice were inoculated with ASF feces purchased from ASF Taconic, Inc. (Hudson, NY), which contained the 8 ASF strains, to established Altered Schaedler Flora (ASF) mice as previously described [30]. Eight-week-old ASF male mice were fed with autoclaved chow and irradiated WSD for 1 week before infecting with 4×10^8^ CFU/per mouse *C. rodentium*, and placed in isolated ventilated caging system (Isocage, Techniplast, Buguggiate VA, Italy) that prevents exogenous bacterial contamination. Fecal samples were collected at indicated days post infection, and the number of *C. rodentium* in feces was counted according to described above. After 35 days, the mice were then removed from isocages and transferred to ABSL2 unit, immediately orally administered 200 μl of the fecal suspension which generated from feces of conventional raised mice, and maintained in a sterile condition (autoclaved cage, food and water).

### STZ treatment and infection

For inducing high glycemia, Eight-week-old male mice fed with GBC were intraperitoneally (I.P.) injected with 40 mg/kg STZ for 4 consecutive days according to previously described [31]. After two weeks, the mice with 5 h fasting glucose between 230 and 265 mg/dl were orally infected with *C. rodentium* at dose of 6 ×10^8^ CFU/per mouse. Mice injected with Sodium citrate buffer (vehicle) was orally infected with the same dose of *C. rodentium* as control. The fecal *C. rodentium* in these mice was measured by using serial dilution and plate counting.

### *C. difficile* cytotoxin assay

Vero cells were grown in DMEM supplemented with 10% FBS (Atlanta Biomedicals), 1% penicillin-streptomycin solution (Corning) at 37°C, and were seeded in 96-well plate to evaluate *C. difficile* cytotoxin as previously described with some modification [32]. Fecal samples were suspended in PBS at 100 mg/ml, and centrifuged at 12000 rpm/min for 10 min to remove the fecal debris and bacteria. Ten-fold serial dilutions of the supernatant was performed, and 10 µl of each dilution were added to the confluent Vero cells which contained 90 µl culture medium. Cell were washed with PBS after 24h incubation, and cell rounding was evaluated, and the cytotoxic titer was expressed as the reciprocal of the highest dilution that caused rounds 100% of Vero cells per gram of sample.

### Cytokine analyses by qRT-PCR

Total RNA was isolated from colon, liver and epididymal fat using Trizol according to the manufacturer’s instructions (Invitrogen). The mRNA expression level of IL-6, α-defense, IL-22 and TNF-α was analyzed by using quantitative RT-PCR (qRT-PCR) according to the Biorad iScript™ One-Step RT-PCR Kit in a CFX96 apparatus (Bio-Rad, Hercules, CA) with the following primers: TNF-α: CGAGTGACAAGCCTGTAGCC, CATGCCGTTGGCCAGGA; α-defensins: GGTGATCATCAGACCCCAGCATCAGT, AAGAGACTAAAACTGAGGAGCAGC; IL6: GTGGCTAAGGACCAAGACC, GGTTTGCCGAGTAGACCTCA; IL-22: GTGCTCAACTTCACCCTGGA, TGGATGTTCTGGTCGTCACC; 36B4: TCCAGGCTTTGGGCATCA; CTTTATTCAGCTGCACATCACTCAGA. Relative expression in transcript levels were calculated by normalization of each amplicon to housekeeping gene 36B4.

### HE Staining

The colons were fixed in 10% phosphate-buffered formalin for at least one weeks at room temperature, before they are transferred into 70% ethanol, embedded in paraffin. Then tissues were sectioned at 5-µm thickness for histological examination by hematoxylin & eosin (H&E) staining. The pathology scores were assessed based on the degrees of hyperplasia and infiltration of immune cells. Crypt length of colon was measured by using image J software.

### ELISA

Fecal samples were homogenized in PBS at 100mg/ml, then centrifuged at 12,000 rpm for 10 min and the supernatant were stored at −80°C. The concentration of LCN-2 in the supernatant was measured by using a DuoSet Mouse Lipocalin-2/NGAL ELISA (R&D Systems) kit. To measure IgA in feces and IgG in serum, 96-well high binding plates were coated with 100µl of bacterial lysate (10 μg/ml in PBS) overnight at 4°C. The plates were washed with PBST (0.05% Tween™ 20 in PBS) and blocked with 1 % BSA for 1 h at room temperature. The fecal supernatant (1:50) and serum (1:1000) were diluted at 1: 50 and 1: 1000 respectively with 1% BSA before applying to 96 well plate. After incubation for 2 h at room temperature, plates were washed and added with goat anti-mouse IgA (SouthernBiotech, 1: 2000) and sheep anti-mouse IgG (GE Healthcare, 1:2000) HRP-conjugated secondary antibody. The optical density (OD) was read at 450 nm (Versamax microplate reader).

### Gut microbiota analysis

Total bacteria DNA was isolated from fecal samples by using QIAamp DNA Stool Mini Kit (Qiagen). To measure the total fecal bacteria number in the feces, quantitative PCR was conducted to analyze the extracted DNA using QuantiFast SYBR Green PCR kit (Biorad) with universal 16S rRNA primers 8F: 5′-AGAGTTTGATCCTGGCTCAG-3′ and 338R: 5′-CTGCTGCCTCCCGTAGGAGT-3′ to measure total bacteria number. Results are expressed as bacteria number / mg feces according to a standard curve. To anayze gut microbiota composition, 16S rRNA gene amplification was conducted as described in the Illumina 16S Metagenomic Sequencing Library preparation guide. Briefly, the extracted DNA was used to amplify the region V4 of 16S rRNA genes by the following forward and reverse primers: 515FB 5′TCGTCGGCAGCGTCAGATGTGTATAAGAGACAGG TGYCAGCMGCCGCGGTAA-3′; 806RB 5′GTCTCGTGGGCTC GGAGATGTGTATAAGAGACAGGGACTACNVGGGTWTCTAAT-3′; These two primers were designed with overhang Illumina adapters. PCR products of each sample were purified with Ampure XP magnetic purification beads (Agencourt) and then run on the bioanalyzer High Sensitivity Chip to verify the size of the amplicon. A second PCR was performed to attach dual indices and Illumina sequencing adapters using Nextera XT Index kit. Products were then quantified before the DNA pool was generated from the purified products in equimolar ratios. The quantity of pooled products was measured before sequencing on the Illumina MiSeq sequencer (paired-end reads, 2 × 250 base pairs) at Georgia Institute of Technology Molecular Evolution Core (Atlanta GA). Following demultiplexing, paired-end reads were quality filtered, denoised, merged and chimera removed using DADA2 plugin in Qiime2. Principal Coordinate Analysis (PCoA) plot based on unweighted UniFrac tables was visualized using Emperor in Qiime2 pipeline. Taxonomy was assigned based on Greengenes 16S rRNA gene database. LEfSe (LDA Effect Size) was used to investigate bacterial members at genus level between groups.

### Immunofluorescence Staining

Mouse colons were embedded in OCT after euthanasia. The tissues were sectioned at 4 μm thickness, and fixed with 4% formaldehyde for 30 min at RT. After washing with PBS, the section was blocked with blocking buffer (Zymed Laboratories Inc., San Diego, USA) before incubated with anti-*Citrobacter* antibody (Abcam, ab37056) overnight at 4°C as previously described[33]. The section was then stained with Secondary Fluorescent Antibody. After washing in PBS, the tissues were counterstained with mounting medium containing DAPI.

### Measurement of ler and tir expression and Rag2/Il2rg DKO infection

The fresh feces samples were collected at 10 days and 26 days post infection, and processed for extraction of total RNAs by using Qiagen RNeasy kit, according to the manufacturer’s protocol. Quantitative real time RT-PCR (qPCR) for ler and tir was performed using a SYBR green PCR master mix and the Step One Real-time PCR system as described previously[9]. Relative expression of ler and tir genes was determined by normalizing to the total *C. rodentium* abundance in feces, which was measured by qPCR using fecal DNA as template. To infect mice, the number of *C. rodentium* in fecal suspension was measured, and a certain amount of fresh fecal suspension including1×10^7^ cfu was orally gavaged into Rag2/Il2rg DKO mice, and the shedding of *C. rodentium* in Rag2/Il2rg DKO mice was monitored after infection.

### Insulin resistance test

WSD fed wild type mice with or without *C. rodentium* persistent infection were maintained on WSD diet for 42 days or 5 months. To conduct insulin tolerance test, mice were fasted for 5 h, and their weight and baseline blood glucose were measured by using a Nova Max plus Glucose meter, then intraperitoneally (I.P.) injected 0.75 U insulin/kg body weight. The blood glucose levels were measured 30, 60, 90 min after injection. After these mice were euthanized, body weight, epididymal fat weight and liver weight were measured.

### *C. rodentium* in vitro culture

Fecal samples were collected from mice fed with WSD and WSD supplemented with inulin, and resuspended in PBS at a concentration of 100 mg/ml overnight at 4°C, then centrifuged at 12000 rpm for 10 min to remove fecal debris and bacteria. 20µl of supernatant was added into 1 ml M9 minimal media (12.8 g/L Sodium phosphate (dibasic) heptahydrate; 6 g/L Monopotassium phosphate; 0.5 g/L Sodium chloride; 1 g/L Ammonium chloride; 0.24g/L Magnesium sulphate, 11.1mg/L Calcium chloride, 100ug/ml kanamycin). This culture medium was inoculated with 1µl LB culture medium containing 1-4×10^6^ CFU *C. rodentium*, and incubated at 37°C with shaking (200rpm). To examine the influence of fat on the growth of *C. rodentium*, 1 ml of M9 minimal media was added with 20 ul of corn oil, or oliver oil (buying from supermarket), or pork lard which was made by heating pork fat to extract oil, then 1-4×10^6^ CFU *C. rodentium* was inoculated, and incubated at 37°C with shaking (200rpm). Growth assay for inulin as carbon source was carried out by adding 20 µl of 20% inulin (Inulin from chicory, Sigma) or 20% D-Glucose as positive control into 1 ml minimal medium, then this culture medium was inoculated with 1µl LB culture medium containing 1-4×10^6^ CFU *C. rodentium*, and incubated at 37°C with shaking (200rpm). WSD and WSD supplemented with inulin were resuspended in PBS at concentration of 100 mg/ml, 20µl of suspension was added into 1 ml M9 minimal culture medium to evaluate its influence on *C. rodentium* growth. To test the influence of short chain fatty acid on *C. rodentium* growth, the minimal cultural medium was supplemented with 10 mM sodium acetate, or sodium butyrate or sodium propionate respectively, 1µl LB culture medium containing 1-4×10^6^ CFU *C. rodentium* was added into the medium, and incubated at 37°C with shaking (200rpm). The growth of *C. rodentium* in the culture medium was measured by using serial dilution and plate counting, expressed as CFU/ml medium.

### Statistical analyses

Statistical significances of results were analyzed by unpaired student t test. Differences between experimental groups were considered significant at *P ≤ 0.05, **P ≤ 0.01 or *** P ≤ 0.001.

## Acknowledgement

We thank Dr. *Etienne-Mesmin Lucie* for help with technical expertise in *C. difficile*. We thank Dr. *Vu L. Ngo* for providing the *C. rodentium* strain.

## Supplemental Figure Legends

**Figure S1.**
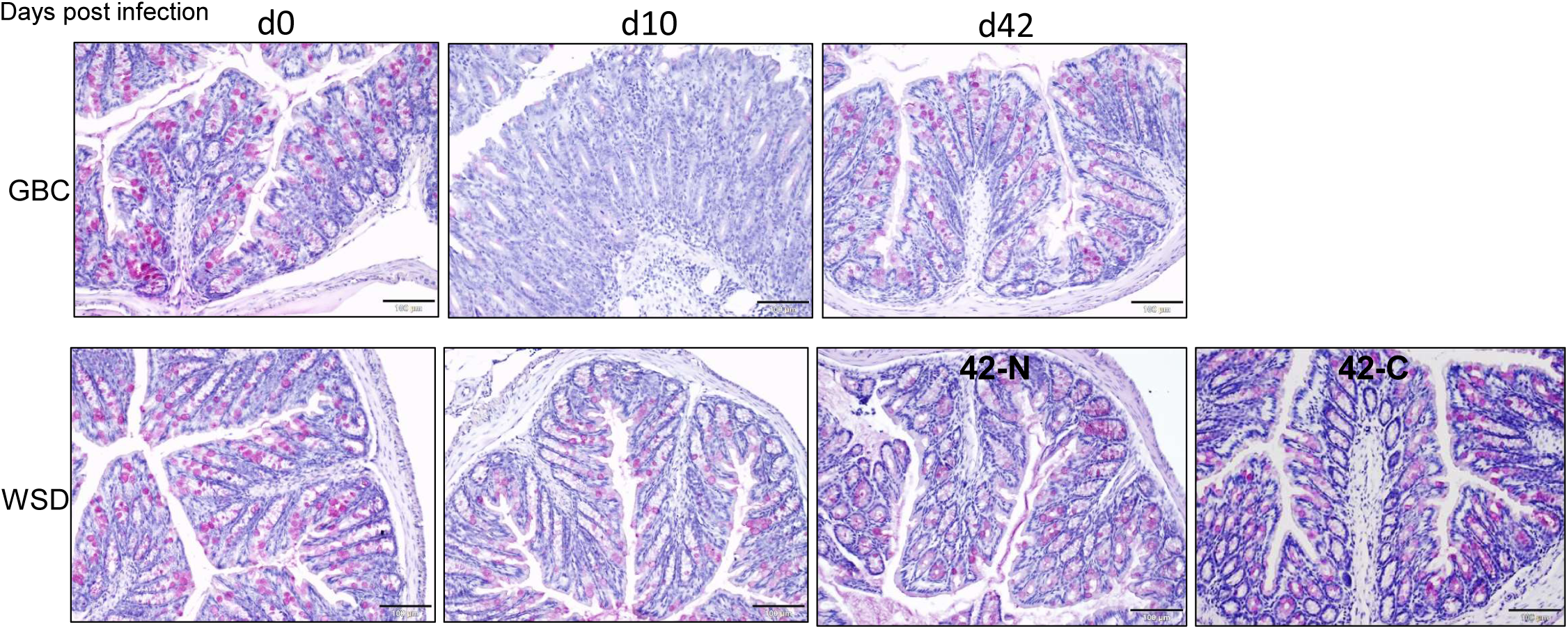
Periodic acid-Schiff staining goblet cells in the crypt of proximal colon. Colon tissue were collected from GBC and WSD fed mice at different time points after infection and stained with Periodic acid–Schiff method.

**Figure S2.**
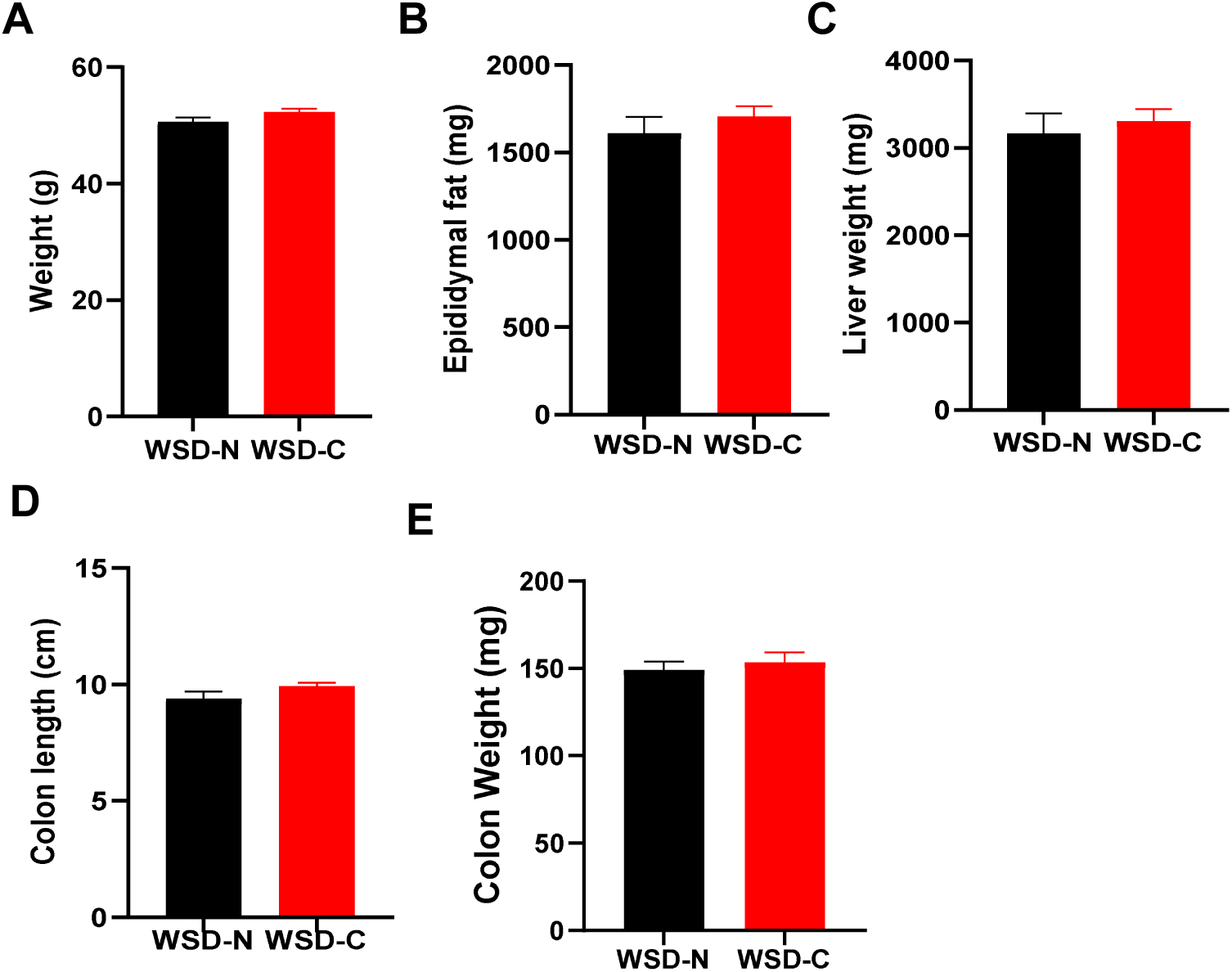
Persistence of *C. rodentium* was not associated with WSD induced obesity. **A-E**. Body weight (A), epididymal weight (B), liver weight (C), colon length (D) and colon weight (E) of WSD fed mice with (WSD-N) or without (WSD-C) *C. rodentium* clearance measured at the end of experiment.

**Figure S3.**
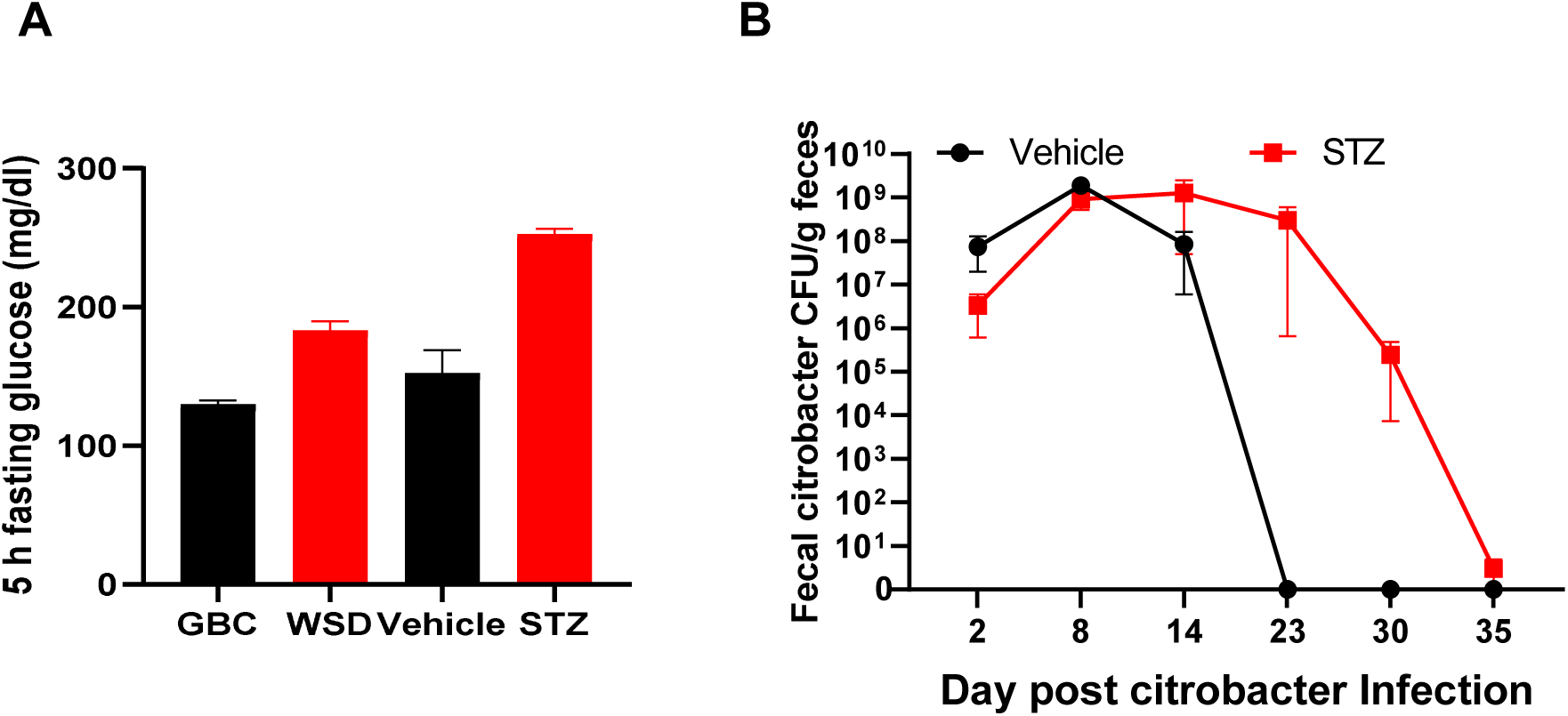
Hyperglycemia delays but does not prevent clearance of *C. rodentium*. **A**. 5 h fasting glucose measured in mice fed with GBC or WSD or treated with STZ. **B**. STZ treated mice with 5 h fasting glucose between 230 and 265 mg/dl were subjected to be infected with *C. rodentium*. The fecal *C. rodentium* was monitored.

**Figure S4.**
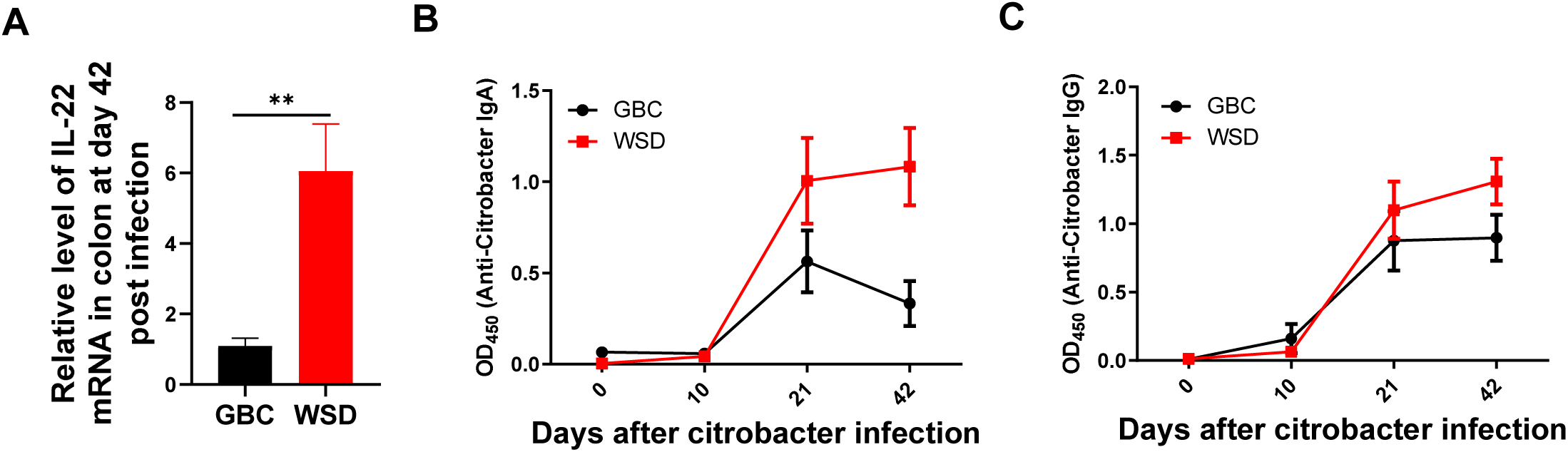
*C. rodentium* persistence despite induced expression of IL-22 and secretion of IgA and IgG. **A**. Relative expression level of IL-22 in colon of mice after 42 days post *C. rodentium* infection. **B-C**. The level of IgA in feces and IgG in serum was measured by ELISA.

**Figure S5.**
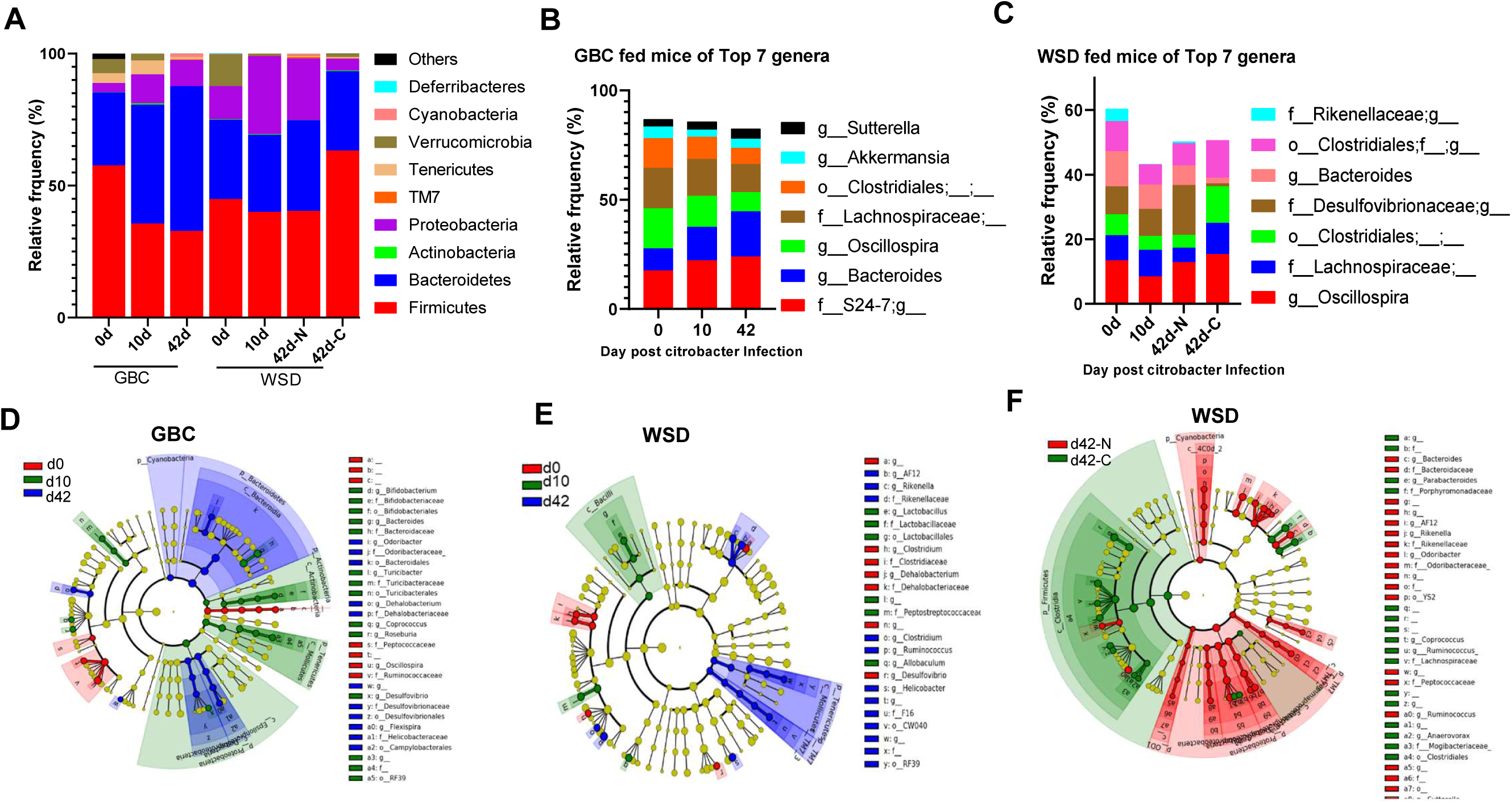
*C. rodentium* infection induces alterations of intestinal microbiota. **A**. Relative abundance of bacteria at the phylum level. **B-C**. The relative abundance of the top 7 genera in GBC (B) and WSD (C) fed mice during *C. rodentium* infection. **D-F**. Cladogram showing different abundant genus in GBC (D) or WSD (E) fed mice during *C. rodentium* infection.

**Figure S6.**
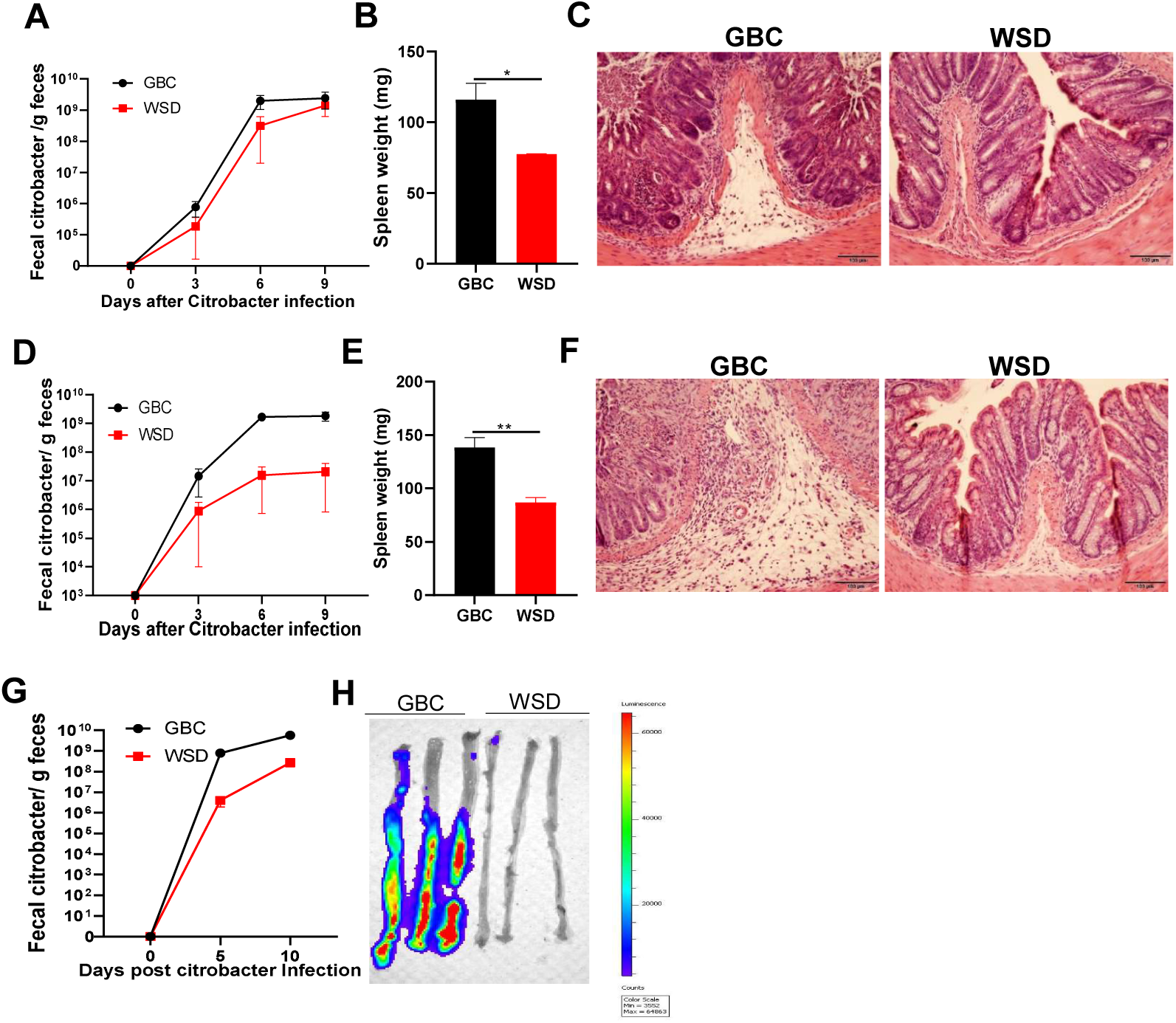
GBC vs. WSD differences in C. rodentium are irrespective of IL-18 and IL-22. IL-18 KO (A-C), IL-22 KO (D-F) and Rag1 KO (G-H) mice were fed with GBC or WSD for 1 week before infected with *C. rodentium*. Fecal *C. rodentium* was measured at indicated days (A, D&G). Spleens were weighted (B&E) and colon was collected to process for HE staining (C&F) for IL-18 KO and IL-22 KO mice; colons collected from Rag1 ko mice were cut longitudinally to remove feces, washed in PBS completely before bioluminescent imaging (H).

**Figure S7.**
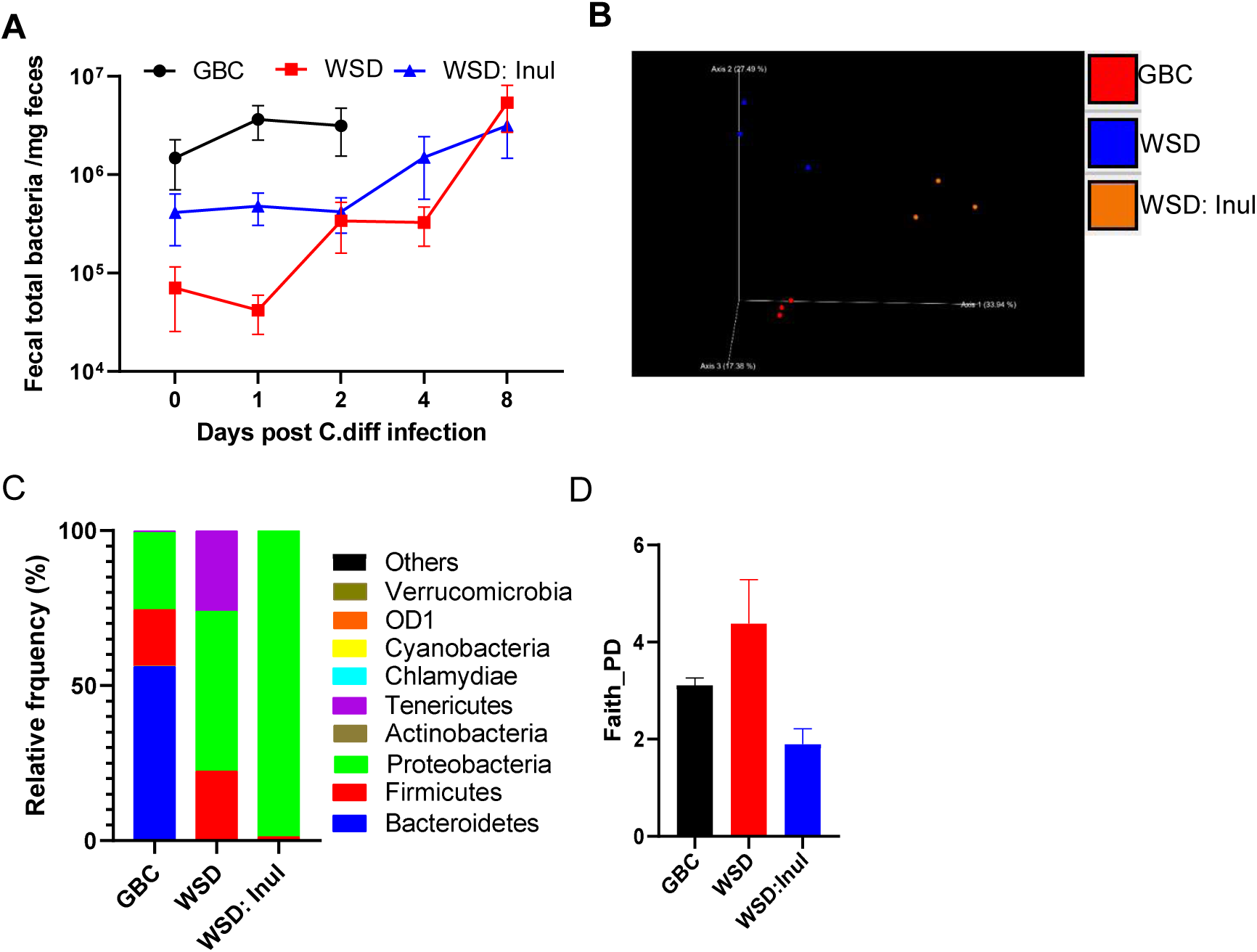
Analysis of gut microbiota during *C. difficile* infection. **A**. qPCR was conducted to analyze the total bacteria in feces during *C. difficile* infection. **B-D**. Fecal microbiota composition was analyzed by 16S RNA sequencing, global composition as expressed by unweighted UniFrac PCoA analysis (B). Relative abundance of bacteria at the phylum level (C). Gut microbiota diversity measured by Faith PD (D).

**Supplementary Table 1.** Diets composition used in this study.

| Product # | D12492 |  | D13081106 |  | D12450J |  |
| --- | --- | --- | --- | --- | --- | --- |
|  | *WSD* |  | *WSD: Inul* |  | *CDD* |  |
|  | gm% | kcal% | gm% | kcal% | gm% | kcal% |
| Protein | 26 | 20 | 23 | 20 | 19 | 20 |
| Carbohydrate | 26 | 20 | 40 | 20 | 67 | 70 |
| Fat | 35 | 60 | 31 | 60 | 4 | 10 |
| Total |  | 100 |  | 100 |  | 100 |
| kcal/gm | 5.2 |  | 4.6 |  | 3.8 |  |
| Ingredient | gm | kcal | gm | kcal | gm | kcal |
| Casein | 200 | 800 | 200 | 800 | 200 | 800 |
| L-Cystine | 3 | 12 | 3 | 12 | 3 | 12 |
| Corn Starch | 0 | 0 | 0 | 0 | 506.2 | 2025 |
| Maltodextrin 10 | 125 | 500 | 75 | 300 | 125 | 500 |
| Sucrose | 68.8 | 275 | 68.8 | 275 | 68.8 | 275 |
| Cellulose, BW200 | 50 | 0 | 0 | 0 | 50 | 0 |
| Inulin, Orafit HP | 0 | 0 | 200 | 200 | 0 | 0 |
| Soybean Oil | 25 | 225 | 25 | 225 | 25 | 225 |
| Lard | 245 | 2205 | 245 | 2205 | 20 | 180 |
| Mineral Mix, S10026 | 10 | 0 | 10 | 0 | 10 | 0 |
| DiCalcium Phosphate | 13 | 0 | 13 | 0 | 13 | 0 |
| Calcium Carbonate | 5.5 | 0 | 5.5 | 0 | 5.5 | 0 |
| Potassium Citrate, 1 H2O | 16.5 | 0 | 16.5 | 0 | 16.5 | 0 |
| Vitamin Mix, V10001 | 10 | 40 | 10 | 40 | 10 | 40 |
| Choline Bitartrate | 2 | 0 | 2 | 0 | 2 | 0 |
| FD&C Yellow Dye #5 | 0 | 0 | 0 | 0 | 0.04 | 0 |
| FD&C Red Dye #40 | 0 | 0 | 0.05 | 0 | 0 | 0 |
| FD&C Blue Dye #1 | 0.05 | 0 | 0 | 0 | 0.01 | 0 |
| Total | 773.85 | 4057 | 873.85 | 4057 | 1055.1 | 4057 |

